# The Evidence Aggregator: AI reasoning applied to rare disease diagnostics

**DOI:** 10.1101/2025.03.10.642480

**Authors:** Hope Twede, Lynn Pais, Samantha Bryen, Emily O’Heir, Greg Smith, Ron Paulsen, Christina A. Austin-Tse, Alex Bloemendal, Cas Simons, Amanda K. Hall, Scott Saponas, Miah Wander, Daniel G. MacArthur, Heidi L. Rehm, Ashley Mae Conard

## Abstract

Variant assessment of rare disease diagnostics depends on using domain knowledge in the time-consuming process of retrieving, reviewing, and synthesizing clinical and technical information. To address these challenges, we developed the Evidence Aggregator (EvAgg), an open-source, generative-AI-based tool designed for rare disease diagnosis that systematically extracts relevant information from the scientific literature for any human gene. EvAgg provides a thorough and current summary of observed genetic variants and their associated clinical features, enabling rapid synthesis of evidence concerning gene-disease relationships. We constructed an expert-curated dataset and evaluated EvAgg’s performance. EvAgg achieves 92% recall in identifying relevant papers, 96% recall in detecting instances of genetic variation within those papers, and ∼80% accuracy in extracting individual case and variant-level content (e.g. zygosity, inheritance, variant type, and phenotype). Further, EvAgg complemented the process of manual literature review by identifying substantial additional relevant information. When tested with analysts in rare disease case analysis, EvAgg reduced review time by 34% (p-value < 0.002) and increased the number of papers, variants, and cases evaluated per unit time. These savings have the potential to reduce diagnostic latency and increase solve rates for challenging rare disease cases.

## Introduction

*Information retrieval* and *synthesis* are fundamental yet challenging tasks, involving the extraction and consolidation of relevant information from vast and often unstructured data sources [1]. These tasks are particularly cumbersome when requiring expertise to sift through scientific literature or knowledge bases [2]. This challenge is core to the rare disease (RD) diagnostic process, where there is often insufficient evidence about variants and gene-disease relationships (GDR) [3], [4].

RDs often have an underlying genetic cause that is challenging to identify. Collectively, RDs affect ∼300 million people worldwide, creating a substantial burden of morbidity and mortality [5] [6] [7]. A genetic diagnosis is critical to accessing reproductive options, treatments, and clinical trials. [3] [5] [7] [8]. While genome and exome sequencing are powerful tools, they identify tens of thousands to millions of variants per patient, with typically only one or two being causal for disease [8].

Bioinformatic filtering can reduce the number of variants requiring human review, but interpretation still involves aggregating information from a variety of sources [9] [10]. This process requires searching for and reviewing academic papers about the disease, genes, and variants in question.

These are time intensive and error prone tasks, making them a major bottleneck for RD diagnosis [9]. Analysts must gather multiple lines of evidence to support or refute a variant’s role in disease, often relying on tools like PubMed [11] or paid resources like Mastermind and Human Gene Mutation Database (HGMD), which aggregate literature-mined variant data [12] [13]. There are publicly funded or community organized curation efforts that provide freely accessible information on GDRs or variant pathogenicity (e.g., OMIM, ClinVar, ClinGen, GenCC, PanelApp) [14] [15] [16] [17]. These resources are critical, but as volunteer-led, retrospective efforts, they struggle to systematically and comprehensively stay up to date with all relevant gene and variant literature and comprehensively capture phenotypic information.

In response to these challenges, researchers have investigated applying natural language processing (NLP) techniques to assist in curation efforts and accelerate diagnostic workflows [12] [18] [19]. Several tools support literature-based evidence gathering in RD case analysis. For example, tools and databases such as Franklin, VarSome, ClinVar, Mastermind, PubTator3, LitVar2, and HGMD index, annotate, or curate literature, helping users locate relevant publications or variant-disease associations [12] [13] [15] [19] [20]. However, these tools are not designed for automated and adaptable extraction of variant-level content and case-level detail (**Supplementary table 1**).

Recently, large language models (LLMs) such as GPT-4 have shown great potential for both information retrieval and complex reasoning tasks [21] [22] [23] [24]. Generative AI (GenAI) has shown promise in other biomedical settings [25] [26]. AI-based approaches have the potential to make previously untenable ongoing literature curation efforts possible [24]. Given their advanced contextual reasoning, we hypothesize that GenAI models like GPT-4 can help variant analysts identify relevant primary literature for case review (Bubeck S, Chandrasekaran V, Eldan R, et al. Sparks of Artificial General Intelligence: Early experiments with GPT-4. *arXiv*; 2023. https://doi.org/10.48550/arXiv.2303.12712) [27].

Variant interpretation platforms such as Franklin and VarSome integrate literature into their workflows, but do not extract or synthesize content [15]. Franklin uses proprietary algorithms and literature references, and its methods are not publicly documented. VarSome cites literature from multiple sources but requires manual content extraction and prioritization. Some ClinVar variant submissions include literature references, but coverage and quality vary by submitter.

Literature search and annotation tools like HGMD, Mastermind, PubTator3, and LitVar2 index, curate, or annotate biomedical literature, though they are not designed for integration into variant interpretation software [12] [13] [19] [20]. HGMD offers manually curated references, with varying time delays, and it requires subscription access. Mastermind indexes full-text articles and supports variant/gene/disease queries but isn’t built to extract or summarize content and requires subscription access. PubTator3 and LitVar2 are open-access tools that use natural language processing for entity recognition and search but are designed for exploration, not structured extraction or synthesis.

Thus, we present the Evidence Aggregator (EvAgg), a modular, open-source tool for traversing the scientific literature while aggregating and synthesizing evidence for any rare disease variant in the human genome (**Figure 1**). Unlike existing tools that focus on indexing or curation, EvAgg ingests papers (e.g. from PubMed), extracts variant-level content, and outputs structured, flexible-format files that can be integrated into any downstream interface. It is designed for speed, transparency, and adaptability, allowing integration of new AI models and customizable verification mechanisms.

**Figure 1:**
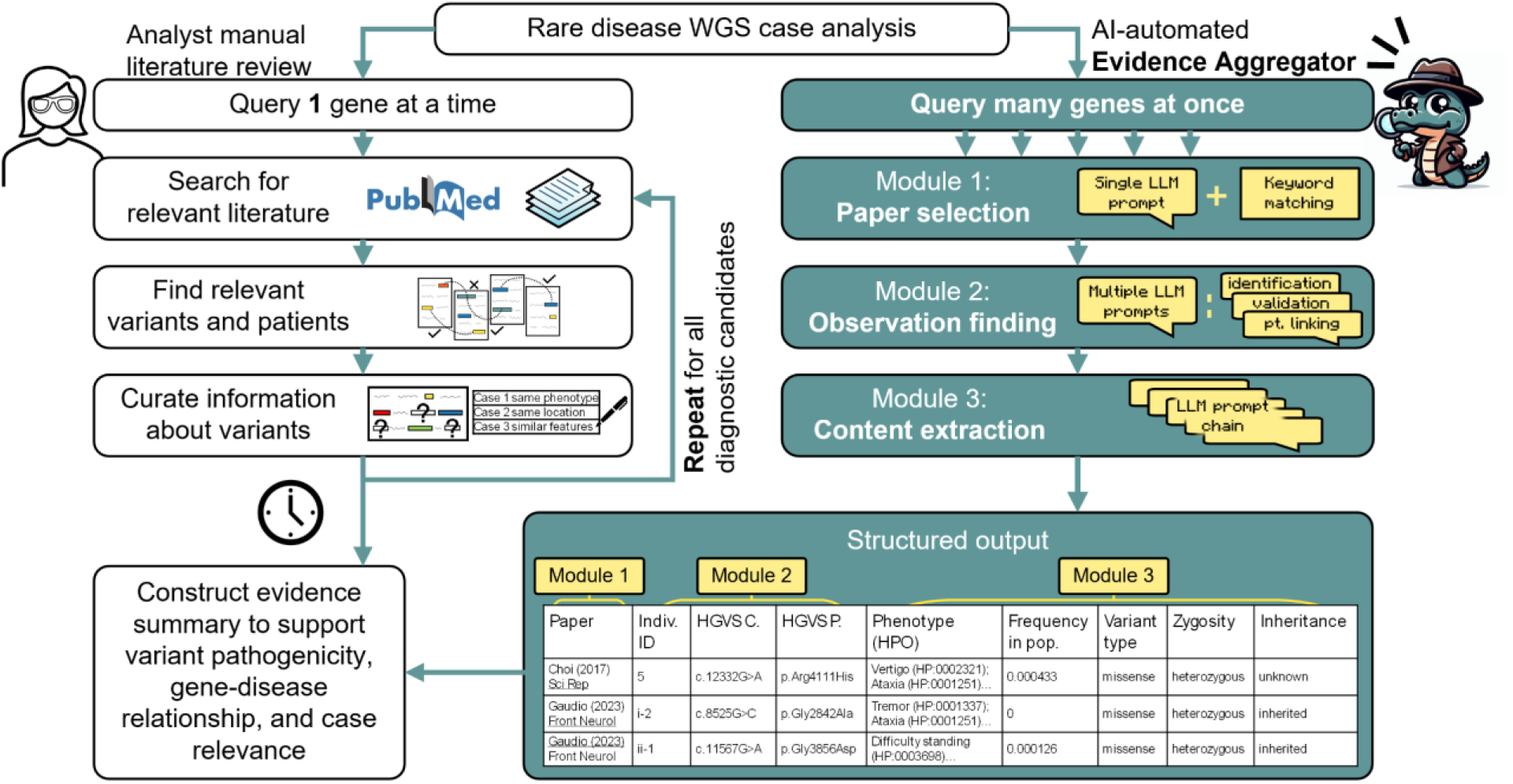
Overview of the Evidence Aggregator. The Evidence Aggregator leverages generative artificial intelligence (AI) to supplement analyst’s literature review tasks, which are currently a manual process in their gene/variant analysis workflow. Analysts conduct literature review for one variant and gene at a time, curating gene-level phenotypes, variant locations, and pathogenicity mechanisms to support or refute variant pathogenicity and relevance to the case under review.

EvAgg complements existing tools by providing an open-source tool for enabling automated, variant-level content extraction and synthesis. It supports traceability and interpretability by flagging uncertain content and logging each step. Unlike manually curated resources, EvAgg can be integrated into reanalysis pipelines to keep variant interpretations current. Compared with similar resources such as HGMD, which provides valuable data elements of variant c. and p. nomenclature, variant type, phenotype, and a publication link, EvAgg delivers these in addition to zygosity, individual-to-variant linkage, population frequency, and variant inheritance.

We evaluated EvAgg’s pipeline performance by comparing its outputs to a manually curated dataset. We also conducted a user study to assess EvAgg’s impact on rare disease case analysis workflows, demonstrating a reduction in case review time as well as an increase in the number of papers, variants, and cases evaluated per unit time.

## Materials and Methods

The core of the EvAgg system is a pipeline that starts with a query gene and through several steps generates structured output describing variant-level observations about a patient, recovered from scientific literature. The larger system can run this pipeline in batches with all the genes for a patient or for a whole cohort of patients. For brevity, this section describes the system as starting with a gene of interest and resulting in evidence for an analyst or clinician to consider during case analysis.

EvAgg’s requirements and evaluation procedure were developed collaboratively by researchers, engineers, and rare disease diagnosticians through an extensive needs-finding exercise [27]. We prioritized recall over precision, as diagnosticians deemed missing relevant content as more harmful than including some extraneous information. For implementation details see **Supplementary Methods**, **Supplementary Figure 1** and our GitHub repository.

### Pipeline overview

The Evidence Aggregator (EvAgg) pipeline is implemented as three independent modules: *paper selection*, *observation finding*, and *content extraction*. Given a query gene, EvAgg first identifies scientific literature describing variants in that gene. EvAgg then extracts all observed variants and individuals (e.g. probands, parents) from those publications, capturing details relevant to variant analysts’ case review (**Figure 1**).

#### Paper selection

EvAgg retrieves candidate publications for each query gene from PubMed Central Open Access (PMC-OA) using REST API calls to the NCBI E-utilities service [28] using the string “`<GENE SYMBOL> pubmed pmc open access[filter]`”. Candidate publications are further filtered to include only those whose licenses permit data mining and derivative use. All subsequent operations are performed on the full text of retrieved publications via the BioC API [29].

EvAgg determines publication relevance using consensus of two methods: keyword matching and LLM-based classification. Keyword matching scans titles and abstracts for inclusion (e.g., “mendelian,” “rare variant”) and exclusion terms to filter out irrelevant content describing non-monogenic disease and somatic cancer. LLM-based classification leverages a single prompt to predict relevance based on the paper’s abstract and title with four few-shot positive and negative examples (see Supplementary Methods). If the methods disagree, the LLM method is run again to break the tie.

#### Observation finding

After identifying candidate publications for each query gene, EvAgg extracts all human-genetic-variant observations related to that gene. This is done using task decomposition with sequential prompts, an established approach for increased LLM performance on complex tasks [30]. First, prompts identify candidate entities: variants, metadata (e.g., genome build transcript IDs), and individual case identifiers. Second, variant strings are normalized to Human Genome Variation Society (HGVS) nomenclature [31], validated, and consolidated so logically linked descriptions (e.g., DNA and protein) are treated as single entities. Given the variability in variant representation [32], EvAgg applies heuristic filters and regular expressions for normalization, then uses Mutalyzer v3 [33] for HGVS syntax and biological validity validation and consequence prediction. Finally, EvAgg links variants to cases to form individual patient-level observations. EvAgg discards papers where it finds no patient-level observations.

#### Content extraction

We identified the categories of content to extract from each observation from a needs-finding exercise with variant analysts [27]. EvAgg uses individual or chained prompts to extract each observation’s phenotype (mapped to HPO terms), zygosity, variant type, and inheritance pattern (see Supplementary Table 2 for value lists). EvAgg also incorporates gnomAD allele frequencies (v4.1) [34] and publication metadata.

#### Presentation of pipeline outputs

EvAgg outputs a JSON formatted file for flexible manipulation and use in downstream systems. As a compelling proof of concept, we have integrated the output as a table in the *seqr platform* [35], a widely used open-source genomic analysis platform designed for rare disease diagnosis and gene discovery. This integration allows EvAgg’s aggregated evidence about a query gene to appear alongside case-specific information.

### Curated dataset

To our knowledge, no ground truth dataset currently exists that provides curated rare disease relevant papers and extracted content across a set of genes. To evaluate EvAgg, we created a publicly available curated dataset through manual curation, the current gold standard, by partnering with expert curators (**Figure 2**). This effort, performed by a subset of the authors (LP, SB, EO, AC, HT, RP, MW; "curation team”), was time intensive and critical for EvAgg’s development and evaluation. We hope this dataset can support the broader community’s efforts (see **Data Availability**).

**Figure 2:**
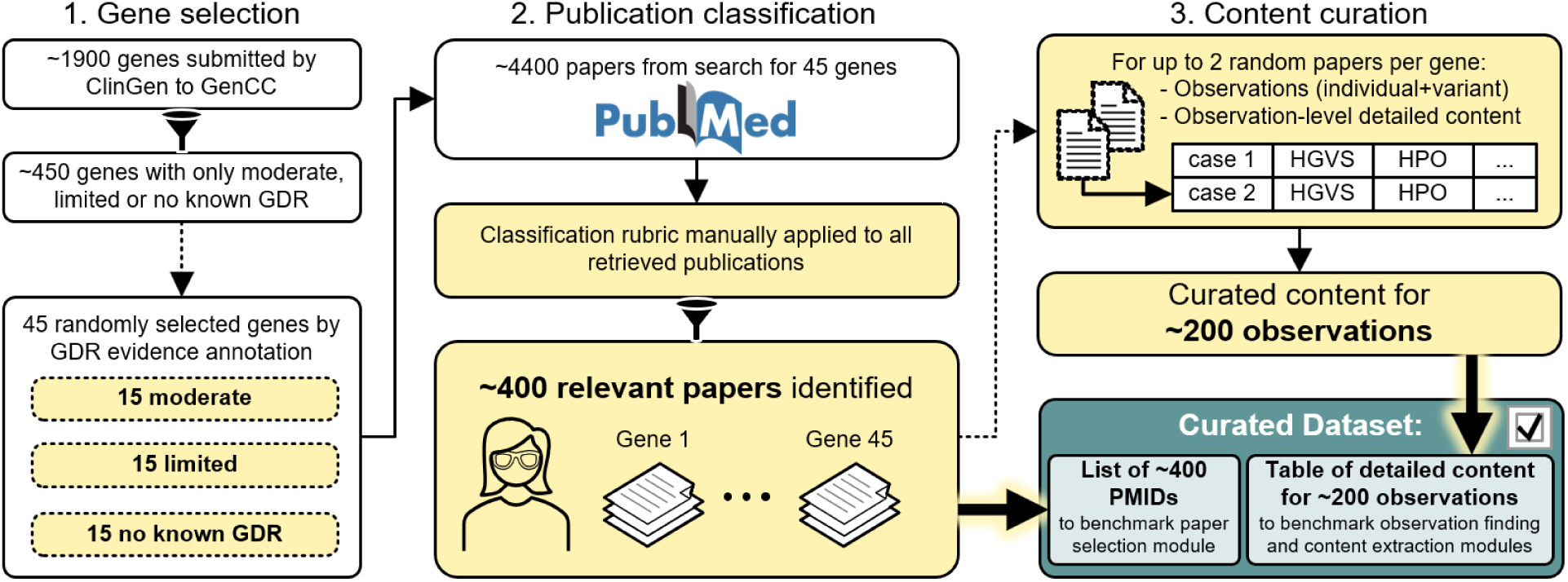
Curated dataset generation process. 1) Forty-five genes were randomly selected from the subset of ClinGen submissions to GenCC with only moderate, limited, or no known gene-disease relationship (GDR). 2) Approximately 4400 papers were returned by a PubMed search for these genes and manually assessed for diagnostic relevance using a standardized rubric, with ∼400 papers deemed relevant (Supplementary methods). 3) From these ∼400 relevant papers, we selected up to 2 papers for each of the 45 genes and curated detailed content for observations of individuals with genetic variants in the gene of interest. The curated dataset, consisting of the complete list of ∼400 relevant papers and curated content for ∼200 observations, was used for subsequent pipeline evaluation. Yellow boxes represent curation of team actions. Funnel-shaped connectors represent filtration steps, dotted line arrows represent random sampling steps, dotted outline shapes represent random samples, bold arrows with yellow highlight represent inclusion of information in the curated dataset. GDR, gene-disease relationship; HGVS, human genome variation society; HPO, human phenotype ontology.

EvAgg is designed to facilitate literature review for genes without well-established GDR. To select genes for the dataset, we began with 1900 genes submitted by ClinGen to GenCC [15] [14], filtered for genes submitted with “Moderate,” “Limited,” or “No known GDR” [14] [15] classifications, and then we randomly selected 15 genes from each category, yielding 45 genes (**Figure 2**).

For each of the 45 genes, our curation team performed a deep review of all papers returned by PubMed search, focusing on publications that appeared after the GDR curation was approved by a ClinGen expert panel. Curators selected relevant papers using a rubric they designed to mirror how variant analysts identify literature with human genetic variant observations that support or refute pathogenicity in RD cases (**Supplementary methods, Curated dataset generation rubric**).

We randomly selected at most two relevant papers for each gene from the PMC-OA subset with licenses that permitted derivative use. We chose only two papers per gene to analyze to limit the amount of work needed to create the dataset. For each of the selected papers, curators extracted individual observation content: publication information, individual case identifiers, variant cDNA and protein HGVS annotation, variant type, variant genome build (e.g. hg19), gene transcript, zygosity, inheritance, phenotype text description, phenotypes encoded as Human Phenotype Ontology (HPO) terms [36], study type, and functional study information when available.

Once curators constructed the curated dataset, we randomly split the curated dataset into a development (dev) and evaluation (eval) set on a per gene basis. 70% of the genes were in the dev set and the remainder in the eval set. The dev set was used for iterative development of EvAgg while the eval set was held in reserve for evaluation after development was completed.

In evaluating EvAgg using the dataset, we also aimed to identify and understand any errors that may have occurred during its construction. After a first evaluation run (see Pipeline evaluation section next), we conducted an in-depth review of putative EvAgg pipeline errors. As part of this review, we sought to identify instances where the EvAgg pipeline was correct, and the error originated from a human mistake when creating the curated dataset. We focused on consistent disagreements (i.e. those recurring across five independent pipeline runs) between EvAgg outputs and the curated dataset. These were formatted as true/false or multiple-choice questions and independently reviewed by two expert variant analysts (LP, SB, EO), blinded to whether responses came from EvAgg or the curated dataset. Disagreements between reviewers were resolved by the broader curation team. Following this error analysis, we updated the curated dataset where warranted. For transparency, evaluation results are reported for both the original and revised datasets.

### Pipeline evaluation

Using our curated dataset, we evaluated the three modules of the EvAgg pipeline: selecting relevant papers, finding observations of human genetic variation within those papers, and extracting specific details about those observations (e.g. an individual’s phenotype).

For paper selection, to determine if EvAgg could find all the papers we were looking for, we measured *recall* as the proportion of curated papers retrieved by EvAgg, and to measure EvAgg’s ability to filter out irrelevant papers we calculated *precision* as the proportion of papers returned by EvAgg that were included in the curated dataset. Observation findings were evaluated similarly, using individual-plus-variant pairs to calculate recall and precision. To distinguish whether discrepancies arose from entity recognition (identifying variants) or entity linking (associating variants with individuals), we separately assessed variant-level performance by treating all individual identifiers as equivalent.

For content extraction, we assessed *accuracy* for all categories (i.e. phenotype, inheritance, variant type, zygosity) by comparing the proportion of exact matches between EvAgg outputs and curated values. Evaluating how well the system extracts phenotypes was more challenging due to the subjectivity involved in mapping free-text descriptions to HPO terms, a task that itself has been the subject of much study [37] [38]. Recognizing that the primary use of phenotypes in rare disease diagnosis is to assess affected organ/systems, we evaluated phenotypes based on matching broader phenotypic categories (**Supplementary methods**).

To evaluate EvAgg’s performance, we adopted a structured evaluation approach with iterative testing on the dev set during pipeline refinement, followed by a final evaluation on the held-out eval set. During development, repeated testing on the dev set helped identify common error modes and guide improvements. After finalizing the pipeline, we ran a single evaluation on the eval set to assess final performance. Each evaluation was run five times, and we report the mean and standard deviation of the resulting metrics. Results are shown for both the original and revised curated datasets.

### User study methods and measurements

We conducted a mixed-methods A/B study to evaluate the EvAgg in a real-world rare disease case review setting (**Figure 3**) We created a simple user interface in the *seqr* variant search platform, where users could access an interactive table of precomputed EvAgg outputs by clicking an “AI Evidence Aggregator” button. Outputs were available for genes with “moderate”, “limited” or “no known” GDR, as described previously. Participants were trained analysts from the participating institutions with prior rare disease case analysis experience within the *seqr* platform.

**Figure 3:**
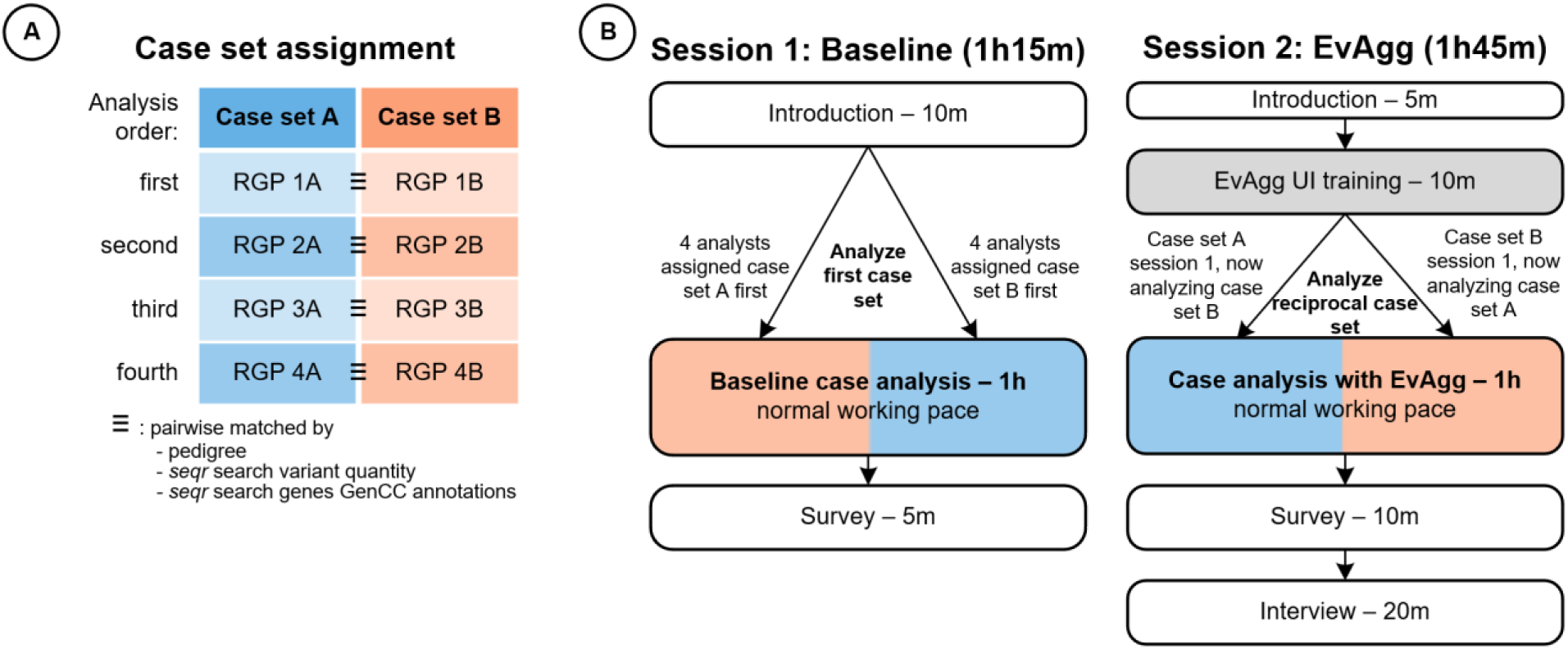
User study design. Participants analyzed unsolved cases from the rare genomes project (RGP) over the course of two days. (A) Cases in each set were presented in a fixed order, with individual cases being pairwise difficulty-matched (i.e. RGP case 1A was of similar difficulty to RGP case 1B). (B) The session 1 (baseline) procedure involved 1 hour of normal case analysis and concluded with a short survey. Participants were randomly assigned one of the two case sets for session 1 and analyzed the reciprocal case set for session 2. During session 2 (EvAgg), participants had EvAgg outputs available. Session 2 concluded with a survey and participant interview. RGP, Rare Genomes Project; ≡, indicates pairwise-matched cases across case sets A and B.

To mitigate time-of-day and fatigue effects, we split the protocol into two virtual, recorded sessions on different days with the same start time. In each session, a study administrator observed a participant analyzing rare disease cases in *seqr* for one hour. The first session served as a baseline; EvAgg was introduced in the second session. During each session, analysts were asked to work at their normal pace and were not expected to complete all cases within the hour, as the intention was to measure effects of EvAgg on their workflow, not to evaluate individual analysts’ performance.

To account for RD case complexity, we created randomized fixed case orders, to approximate difficulty-matched sets of four unsolved cases selected from the Broad Institute’s Rare Genomes Project (raregenomes.org). Individual cases were approximately matched by affected status of family members in pedigree, quantity of variants returned in *seqr* variant search, and ClinGen/GenCC annotation of genes returned in *seqr* search. Participants were randomized to one of the case sets on the first day, analyzing the reciprocal set on the second day (**Figure 3**).

After using EvAgg, participants completed Likert-scale surveys assessing system usability, trust, and overall experience [39] [40]. We conducted semi-structured interviews with participants exploring specific features influencing sentiment and trust, importance of accuracy, and how EvAgg may impact their workflow. An inductive approach for emerging themes was used to analyze qualitative data from participants’ responses to the survey and interview questions.

We measured workflow changes from time-stamped user actions and artifacts from *seqr*. We used a mixed-effects model to analyze the impact of EvAgg on individual behavioral metrics, accounting for both fixed effects (study session and case set) and random effects (individual participants).

## Results

We evaluated the Evidence Aggregator (EvAgg) against manual curation by expert analysts. This evaluation reflects EvAgg’s unique role as an automated, variant-level literature aggregation and synthesis tool - capabilities not unified in existing tools.

### Curated dataset characteristics

We built the curated dataset from 4401 PubMed search results. Literature volume varied widely by gene, with mean M=112.85, standard deviation SD=92.65 papers per gene reviewed. Of these, curators identified M=10.05, SD=15.11 papers per gene as relevant (392 total), meaning only 1 in 11 papers contained case or variant-level evidence useful for gene-disease evidence review. Of the set of 392 relevant papers, 133 met the licensing requirements for access and appeared in the PubMed Open Access subset. Among these, 41 papers (covering 24 of 45 genes) contained variant observations. Curators extracted 224 unique observations representing 167 different variants. On average, each gene had M=9.33, SD=12.91 case observations (M=6.96, SD=10.09 unique variants) and each paper had M=5.60, SD=9.53 case observations (M=4.28, SD=8.13 unique variants).

### Error analysis

Despite strong overall performance, EvAgg showed 470 consistent disagreements with the initial curated dataset across paper selection, observation finding, and content extraction. Of these, 74 (15.7%) involved paper selection, 81 (17.23%) observation finding, and 226 (48.1%) phenotypic feature extractions from 52 distinct observations.

Manual review determined that EvAgg provided more relevant or accurate outputs than the original curated dataset in 235 (50%) cases. These findings highlight the complexity of manual literature curation and the potential for AI tools like EvAgg to enhance information retrieval process (see **Figure 4C**, **Supplementary table 6**).

**Figure 4:**
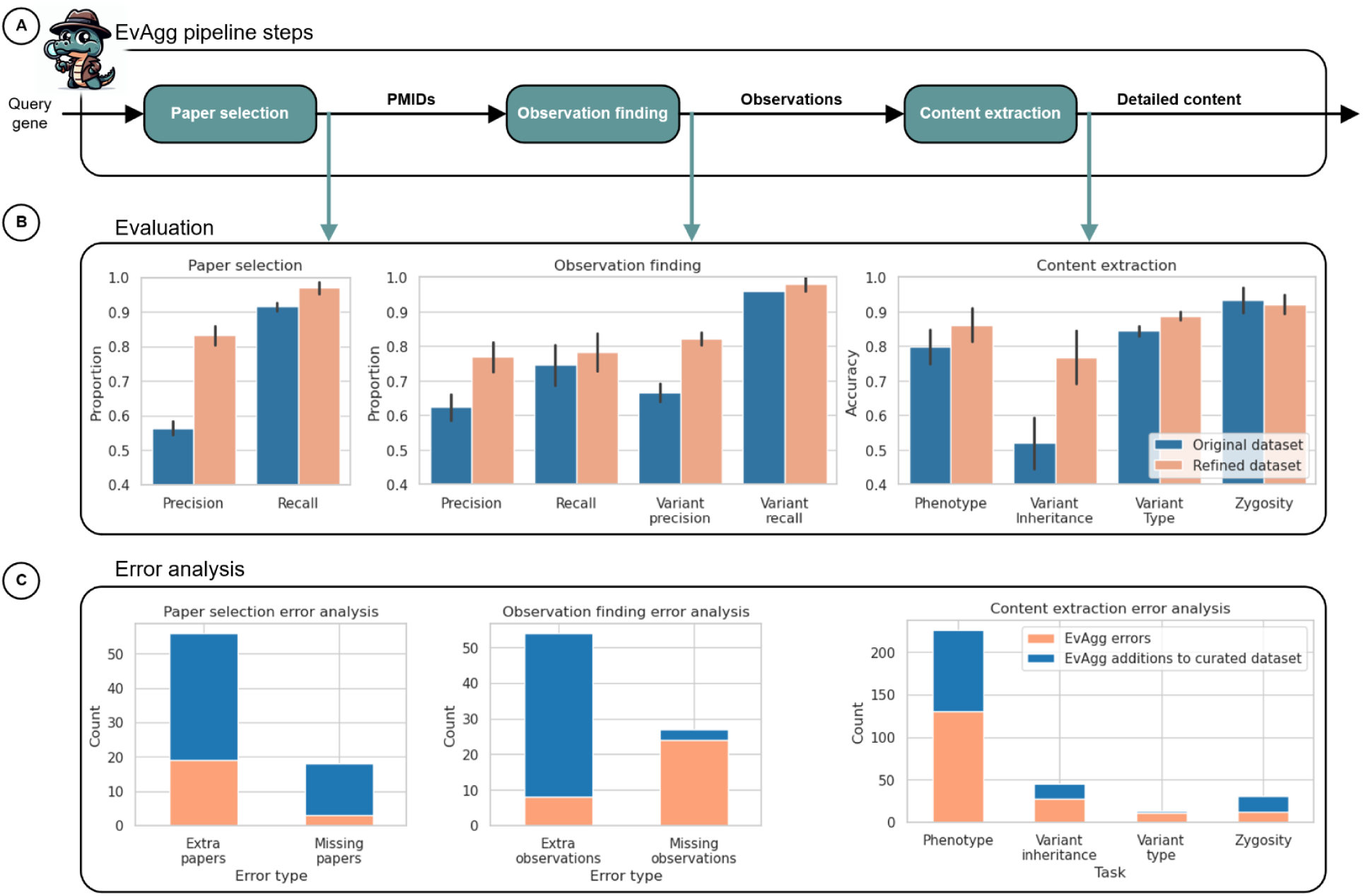
Pipeline evaluation overview and results. (A) Individual steps of the EvAgg pipeline were separately evaluated against both the original and a refined version of the curated dataset to assess tool performance in different aspects of the task. (B) Task performance was evaluated for the two versions of the curated dataset: precision and recall for the paper selection and observation finding tasks, accuracy (i.e., the frequency of matching curated dataset content) for the content extraction task. (C) Breakdown of error analysis shows EvAgg errors (orange) and EvAgg additions to the refined version of the curated dataset (blue). The Evidence Aggregator character icon was AI-generated using DALLE·3.

### Pipeline performance

The following results discuss performance using the evaluation subset of our curated dataset, assessed before and after curated dataset revision. Unless noted, all evaluation metrics refer to the pre-revision version of the dataset. **Supplementary results** include full evaluation analyses (devevelopment and evalaluation, pre- and post-revision) and a performance comparison of GPT-4- Turbo with other candidate LLMs as the underlying LLM used by the system.

### Selecting relevant papers

EvAgg consistently selected similar sets of papers, with each paper appearing in ∼89.3% of runs, showing stable performance despite LLM variability. It achieved high recall, retrieving M=0.92, SD=0.012 of curated relevant papers (**Figure 4b**), averaging 3.2 false negatives (out of 38). Although we prioritized recall during development, precision increased substantially after error analysis, from M=0.56, SD=0.019 to M=0.83, SD=0.028, because reviewers reclassified 37 of 56 original false positives as relevant.

### Finding observations of genetic variation

We evaluated EvAgg’s ability to identify specific variant observations in curated papers. It achieved M=0.75, SD=0.058 recall for observation finding. The system did much better at recall for identifying relevant variants regardless of individual association reaching M=0.96, SD=0.0. This difference reflects challenges in correctly identifying individuals in a paper or correctly linking individuals to the variants.

### Extracting detail about observations

EvAgg achieved an average phenotype classification accuracy of M=0.80, SD=0.050; zygosity M=0.93, SD=0.036; variant type M=0.84, SD=0.014; and variant inheritance M=0.52, SD=0.073 (**Figure 4b)**. A single paper accounted for 16 of 45 discrepancies between the original and revised curated datasets. Error analysis led to updates for 12 of those discrepancies, explaining the performance shift.

Generalized phenotype concordance between EvAgg and the curated dataset reached M=0.86, SD= 0.047. On average, EvAgg reported more phenotype terms than the curated dataset (M=3.9, SD=4.0 vs. M=2.7, SD=3.1, resulting in high recall (M=0.96, SD=0.025) and slightly lower precision (M=0.89, SD=0.033) (**Supplementary table 8, Supplementary figure 3**). These results suggest that EvAgg reliably captures almost all relevant phenotypes, with most errors involving extra, not missing terms.

### User study in rare disease case analysis

We enrolled eight highly trained participants with 1-10 years of experience in rare disease genome case analysis. We conducted the study within collaborating institutions and implemented it within the *seqr* platform. Two participants contributed to curated dataset generation but did not participate in study design, development, execution, or analysis, and had no exposure to EvAgg in *seqr*. We excluded these two participants from survey and interview feedback from analysis. We collected detailed notes and tracked objective workflow metrics: number of EvAgg interactions, cases/variants reviewed, publications read (opened full-text and reviewed), and time spent per case/variant. With EvAgg, participants reviewed cases 34% faster M=27.2, SD=11.3 vs M=17.9, SD=4.8 min/case, p<0.002) and reviewed more papers (M=2.5, SD=2.2 vs M=6.0, SD=4.8, p<0.02) without increasing manual publication searches (p=0.39) (**Figure 5**).

**Figure 5:**
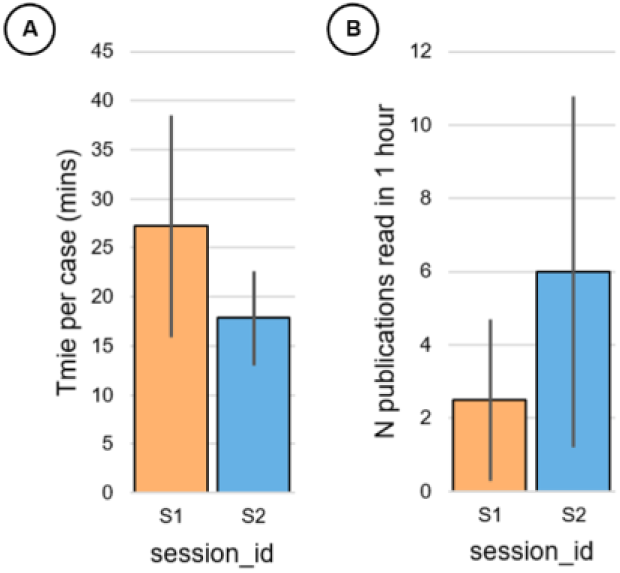
Workflow changes. Changes measured from user study in terms of (A) time per completed case analysis and (B) total number of publications read during 1 hour of case analysis. S1, measure from user study session 1 (baseline without EvAgg, orange bar); S2, measure from user study session 2 (case analysis with EvAgg available, blue bar).

All participants reported positive sentiment and believed they could learn to use EvAgg quickly. All participants perceived time savings, though some noted that “rabbit holing” or inexperience could reduce efficiency. Half said they would shift focus to other tasks, and most also said they would use the saved time to explore cases more deeply. Participants primarily used EvAgg early in their review process.

Participants indicated mixed trust and expressed they would feel more confident after understanding how EvAgg works and its limitations. While all emphasized the importance of accuracy (“*it’s patients that we’re assessing”*), they prioritized accuracy in phenotype and variant location/change. Every participant tolerated some inaccuracy, noting they would double check all information and expect minor errors (“*I’m not taking the information in the table blindly*”).

## Discussion

Our study demonstrates that GenAI tools have the potential to significantly improve information retrieval and reasoning in rare disease diagnostic analysis, reducing the time and effort required for analysts. EvAgg effectively retrieves and synthesizes relevant literature, achieving high sensitivity, specificity, and accuracy in pipeline evaluations. EvAgg recalled over 95% of relevant rare disease papers and variants, accurately linked variants to individuals, and extracted key content. Through error analysis, we found that EvAgg identified additional relevant papers, observations, and content missed during manual curation, highlighting GenAI’s value in complex, domain-specific tasks.

Our user study shows that experienced analysts used EvAgg to analyze cases more efficiently. Participants completed more cases, reviewed more variants, and read more papers per case. They expressed strong interest in integrating EvAgg into their workflows, describing case analysis as easier and requesting broader gene coverage. Participants emphasized the importance of understanding EvAgg’s performance and error modes to build trust.

Our rigorous pipeline evaluation and curated dataset support accurate performance metrics, which contribute to user confidence. We have published the curated dataset on GitHub and welcome feedback and contributions.

GenAI tools like EvAgg offer a scalable accelerator of manual curation and custom NLP pipelines, which require extensive training data and compute. By making summarized information more accessible, GenAI can democratize knowledge and deliver timely insights to researchers and clinicians.

In rare disease diagnosis, where time is limited and literature review is demanding, GenAI tools like EvAgg can ease the burden and uncover overlooked insights. By releasing EvAgg and its pipeline evaluation dataset openly, we invite the community to evaluate, improve, and integrate this tool -advancing a shared goal of faster, more informed rare disease diagnosis.

## Limitations

EvAgg’s performance depends on the underlying model. Future models may outperform GPT-4- Turbo, which we used in most pipeline evaluation analyses. High model costs may limit adoption in resource-constrained settings. To improve future versions, we plan to explore strategies like text chunking and few-shot examples. Currently, EvAgg supports only English inputs and outputs.

Although we followed a rigorous process to develop the curated dataset, its small size highlights the need for larger, shared datasets and pipeline evaluations that include more analysts and cases. On the tool usability front, our user study revealed workflow changes, but larger, longitudinal studies will be required to quantify EvAgg’s impact on user experience during case analysis.

EvAgg was designed to provide specific information and is thus not exhaustive. While EvAgg complements existing tooling, a direct comparison with most analogous tools could not be made due to license restrictions. We did not evaluate performance or cost for genes with strong or definitive evidence for disease relationships, which would typically have more extensive literature. We also did not process abstracts or user-provided files for closed-access papers. Expanding open access and permissive licensing remains essential to improve GenAI and other computational tools for literature synthesis.

## Data and code availability

The Evidence Aggregator code is fully available and documented in an open-source GitHub repo: https://github.com/cassimons/healthfutures-evagg. Datasets generated and/or analyzed here are available in the GitHub repo data/ directory (curated dataset, user study quantitative data, error analysis results) and in supplementary information (user study qualitative data). Scripts to regenerate manuscript quantitative findings are available in the Github repo scripts/ directory.

## Author contributions

All authors were involved in the development of the research proposal. AMC, HT, LP, SB, EO, RP, and MW created the curated dataset and performed error analysis comparing the curated dataset to pipeline outputs. AMC, MW, and GS developed the EvAgg software package. HT, AMC, and AKH designed user study. HT executed the user study. AMC, HT, and MW performed data analysis. HT, RP, AMC, and MW conducted responsible AI review. HT, AMC, GS, RP, SS, MW, HR, CAA-T, LP, EO, AB, CS, SB, and DM were involved in the authoring of the manuscript.

## Funding statement

The authors declare the following competing interests: HT, AMC, GS, RP, AKH, SS, and MW are all full-time employees of Microsoft. HR, CAA-T, LP, EO, AB, CS, SB, and DM receive research funding from Microsoft.

## Ethics declaration

User study design underwent IRB review by Microsoft Research IRB#1 (OHRP-IRB00009672, IORG0008066). Informed consent was obtained from all participants as required by the IRB. Participant data was de-identified for analysis and distribution.

## Declaration of AI and AI-assisted technologies in the writing process

Statement: During the preparation of this work the author(s) used Copilot (Microsoft Bing Chat) to reduce abstract length and DALLE•3 image creator to generate the stylized cartoon “Evidence Aggre-Gator” character/logo. After using these tools/services, the authors reviewed and edited the content as needed and take full responsibility for the content of the publication.

## Notes

https://github.com/cassimons/healthfutures-evagg

### Competing Interest Statement

The authors declare the following competing interests: Hope Twede, Ashley Mae Conard, Greg Smith, Ron Paulsen, Scott Saponas, Amanda K. Hall, and Miah Wander are all full-time employees of Microsoft. Heidi Rehm, Christina A. Austin-Tse, Lynn Pais, Emily O'Heir, Alex Bloemendal, Cas Simons, Samantha Bryen, and Daniel MacArthur receive research funding from Microsoft.

### Summary of Updates

a reduction in word count standardization of terminology used throughout the paper simplified language about the user study design display of pre- and post-error analysis performance evaluation results revision of user study qualitative analysis to remove authors figure modifications and some shifts to supplement addition of a table comparing features to other tools shifting user study thematic analysis data to a table

