## Supplementary Information for "The Evidence Aggregator: AI reasoning applied to rare disease diagnostics"

##### Supplementary introduction

| Feature / Tool | Franklin | VarSome | ClinVar | HGMD | Mastermind | LitVar2 | PubTator3 | EvAgg |
|---|---|---|---|---|---|---|---|---|
| Variant-level literature linkage | + | + | + | + | + | + | + | + |
| Relevant content extraction | – | – | – | – | – | – | – | + |
| Automated synthesis | – | – | – | – | – | – | – | + |
| Open-access | – | –* | + | – | –* | + | + | + |
| Open-source | – | – | + | – | – | + | + | + |
| Customizable AI model integration | – | – | – | – | – | – | – | + |
| Transparent extraction process | – | – | – | – | – | – | – | + |
| Verification of extracted content | – | – | – | – | – | – | – | + |
| Flagging uncertain content | – | – | – | – | – | – | – | + |

| | | | | | | | | |
|---|---|---|---|---|---|---|---|---|
| Flexible output for integration | – | – | – | – | – | – | – | + |
| Supports reanalysis | – | – | – | – | – | – | – | + |

*VarSome and Mastermind offer limited free access but require subscriptions for full functionality.

***Supplementary table 1: EvAgg feature/tool comparison.*** *EvAgg features as compared to popular open-access or commercially available variant-level literature curation resources.*

### Supplementary methods

#### Supplementary pipeline overview and design criteria information

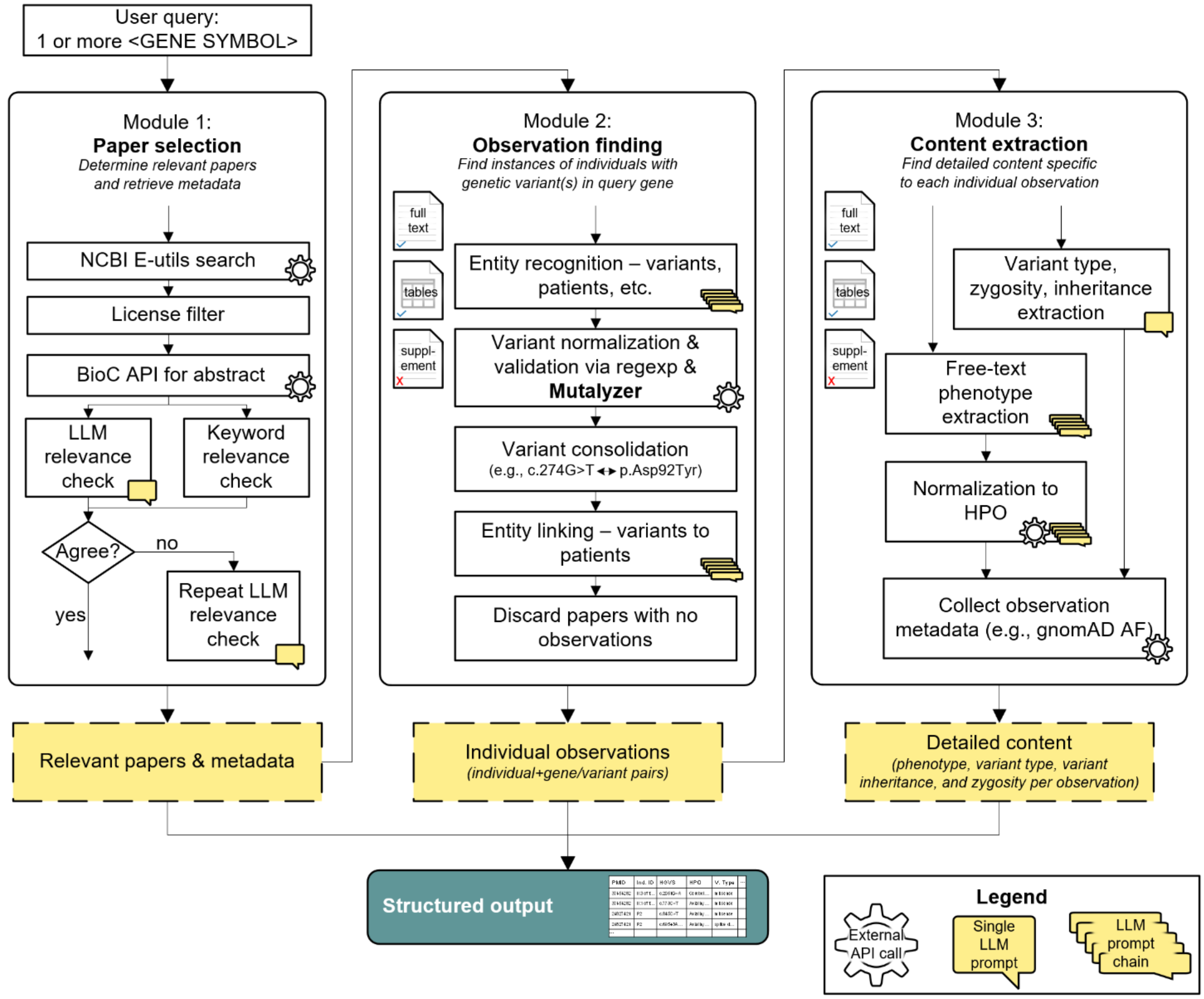


***Supplementary figure 1: EvAgg schematic.*** *This figure depicts the processing steps that take place within each module of the EvAgg tool. The first module is responsible for identifying PubMed papers that are potentially diagnostically relevant for the gene of interest. The second module has the task of identifying observations of human genetic*

*variation in those papers, where a unique observation is specified by a gene, HGVS variant description, and individual case identifier. The third module then extracts specific details about those individual observations, such as phenotypes and inheritance patterns. EvAgg uses a combination of internal logic, API calls to external web resources, and prompts or prompt chains issued to the LLM. Dashed-outline yellow boxes represent content included in structured output. NCBI, National Center for Biotechnology Information; LLM, large language model; regexp, regular expression (text pattern matching); HPO, Human Phenotype Ontology; AF, allele frequency; API, application programming interface.*

##### *Paper selection*

In addition to parametrizing paper search based on the query gene, the tool can be configured to limit the search of PMC-OA based on publication date and the maximum number of papers to return. This was leveraged for analytical repeatability and to facilitate comparison with the curated dataset.

Papers describing only structural variants were considered out of scope for EvAgg at this time and were filtered out during the publication relevance checking stage.

LLM-based classification few-shot positive and negative examples were drawn from the development subset of the curated dataset data ensuring that no examples drawn from a particular query gene were used when that gene was being run through the pipeline.

##### *Observation finding*

The process to identify all observations of human genetic variation associated with the query gene from selected publications was complicated by variability in how genetic variation is expressed in the literature, ambiguity in language linking genetic variants to the individuals who possess them, fragmentation of related information throughout the text, and accidental typographic errors. As a result, this pipeline module included a series of *entity recognition* (i.e., finding the correct variants within a paper), *entity linking* (i.e. correctly associating those variants with individuals), and *fact-checking* (i.e. verifying the factual accuracy) stages, full discussion of which is outside the scope of this manuscript.

Note that a given variant could be involved in multiple observations in a paper, as was the case in studies of an extended family pedigree with an inherited pathogenic variant. Or a given individual can possess multiple variants, as was the case in individual cases with compound heterozygosity.

##### *Content extraction*

See **Supplementary table 2** below for Module 3 (content extraction) values.

| **Extracted content category** | **Possible EvAgg output strings** |
|---|---|
| **Variant type** | "missense", "frameshift", "stop gained", "splice donor", "splice acceptor", "splice region", "start lost", "inframe deletion", "frameshift deletion", "inframe insertion", "frameshift insertion", "structural", "synonymous", "intron", "5' UTR", "3'UTR", "non-coding", "unknown" |
| **Variant inheritance** | "inherited", "de novo", "unknown" |
| **Variant zygosity** | "homozygous", "heterozygous", "compound heterozygous", "none" |

| | |
|---|---|
| **Phenotype** | Only required to be a valid entity in the HPO; all curated dataset terms are children of the following generalized organ/system description terms used for pipeline evaluation:<br>HP:0040064 \| Abnormality of limbs<br>HP:0000769 \| Abnormality of the breast<br>HP:0025354 \| Abnormal cellular phenotype<br>HP:0001939 \| Abnormality of metabolism/homeostasis<br>HP:0000152 \| Abnormality of head or neck<br>HP:0002715 \| Abnormality of the immune system<br>HP:0001574 \| Abnormality of the integument<br>HP:0002086 \| Abnormality of the respiratory system<br>HP:0001197 \| Abnormality of prenatal development or birth<br>HP:0000818 \| Abnormality of the endocrine system<br>HP:0025142 \| Constitutional symptom<br>HP:0000707 \| Abnormality of the nervous system<br>HP:0025031 \| Abnormality of the digestive system<br>HP:0001608 \| Abnormality of the voice<br>HP:0001871 \| Abnormality of blood and blood-forming tissues<br>HP:0000598 \| Abnormality of the ear<br>HP:0001626 \| Abnormality of the cardiovascular system<br>HP:0000478 \| Abnormality of the eye<br>HP:0045027 \| Abnormality of the thoracic cavity<br>HP:0001507 \| Growth abnormality<br>HP:0033127 \| Abnormality of the musculoskeletal system<br>HP:0002664 \| Neoplasm<br>HP:0000119 \| Abnormality of the genitourinary system |

***Supplementary table 2: EvAgg's pipeline outputs for evaluating content extraction task performance***. *EvAgg Module 3 (content extraction) outputs for variant type, variant inheritance, and zygosity are constrained to return a single value from each category's set of text string values (separated by commas). Zero or more individual phenotype HPO terms may be extracted for each observation. Each individual phenotype HPO term is only constrained in that it must be a valid entity from the HPO. In the case that the phenotype extraction prompt chains cannot successfully determine a valid HPO entity corresponding to a free text phenotype, the original free text is returned by the pipeline, without a corresponding HPO term. See supplementary pipeline evaluation methods for additional information about phenotype pipeline evaluation.*

For each observation, EvAgg retrieved many additional pieces of information (content) from the source text and other databases. Population frequency values are as reported in gnomad v4 maximum credible genetic ancestry group allele frequency. In addition to the content categories described in the main text, the pipeline can be configured to extract study type for the paper being reviewed and any follow-up functional studies that were performed. However, discussion of these categories is outside the scope of this manuscript.

#### Prompt implementation detail

When issuing prompts to the LLM, EvAgg provides the entirety of the text of each paper to the model. In addition, when searching for variants, individuals, or phenotypes, EvAgg will also run the associated prompts separately on any tables within a paper, as these are often enriched for the entities of interest. EvAgg retains the union of results from applying these prompts to the full text and the individual tables.

Due to this design decision, we restricted model selection to Azure OpenAI (AOAI) Service models supporting context sizes of 128k+ tokens available at the time of development and evaluation: GPT-4-Turbo, GPT-4o, and GPT-4o-mini.

#### Paper selection prompt implementation details

##### *Prompt-optimization*

With the advent of the reasoning capabilities of large language models, we devised a series of prompting techniques to enhance the task of finding relevant papers. Initially, we created a generic prompt based on rubric information and implemented a paper-finding prompt optimization schema to systematically refine the prompt text by adjusting the language and running pipeline evaluation to assess overall performance improvement.

Specifically, the schema follows these steps: first, it runs the paper-finding pipeline with the original prompts. Second, it executes the paper-finding pipeline evaluation. Third, if the precision and recall are not both 1.0, it utilizes the LLM prompt assistant to modify the prompt in hopes of improving its effectiveness. Fourth, it reruns the paper-finding pipeline and pipeline evaluation. Finally, it checks if the precision and recall have improved by more than a specified threshold (epsilon) after a minimum number of iterations and continues until a maximum number of iterations is reached.

##### *Few-shot learning*

Few-shot learning has been extensively shown to enhance the effectiveness of prompt engineering [41]. For our task of relevant paper selection, we compared various combinations of three common approaches: chain-of-thought, few-shot positive and negative examples (title and abstract), and few-shot positive and negative examples (full text). Our results (not shown, considered out of scope for this publication) indicated that the best performance was achieved using few-shot learning with positive and negative examples based solely on the title and abstract.

Our methodology involved several steps. Initially, we gathered all papers in the development set that did not include the query gene. These papers were then clustered using three common approaches: Elbow, Silhouette, and Bayesian Information Criterion, resulting in four predominant clusters. Within each cluster, we randomly sampled a gene and a paper (ensuring each gene was selected only once across clusters) and appended that paper's title and abstract to the end of the prompt.

Next, we ran EvAgg using the few-shot positive example methodology, followed by the associated pipeline evaluation to identify irrelevant papers. These irrelevant papers were then clustered and sampled following the same steps as the positive examples to generate a set of four negative few-shot examples. The positive and negative examples were derived from both the EvAgg output and a curated dataset, neither of which are perfect as there is no true ground truth. Although unlikely, there is a possibility that if both EvAgg and the curated dataset incorrectly classified a relevant paper as irrelevant, and it was subsequently included in the negative examples, this would result in a false negative. Finally, we ran the EvAgg again with the few-shot positive example methodology, followed by the pipeline evaluation.

#### Curated dataset generation

The final set of 45 genes used for curated dataset generation were derived from the subset genes where the most recent ClinGen-submitted GDR to GenCC was "Moderate" (n=205), "Limited" (n=220), or "No known GDR" (n=16), with first ClinGen GenCC submission date for these genes recorded as between 1976 and 2023 depending on the gene (see **Supplementary table 3** and **Supplementary table 4** below for gene list and publication detail) [14] [15].

| Set | Gene | Evidence Base | Total Papers | Accessible Proportion | # PubMed Papers | GenCC ClinGen submission |
|---|---|---|---|---|---|---|
| **dev** | GRXCR2 | Moderate | 6 | 83 | 9 | 2014 |
| | B4GAT1 | Moderate | 4 | 50 | 19 | 2013 |
| | COG4 | Moderate | 18 | 44 | 53 | 2009 |
| | TOP2B | Moderate | 8 | 38 | 92 | 2019 |
| | EMC1 | Moderate | 9 | 33 | 27 | 2016 |
| | MLH3 | Moderate | 33 | 30 | 263 | 2001 |
| | SLFN14 | Moderate | 11 | 27 | 18 | 2015 |
| | JPH2 | Moderate | 28 | 25 | 107 | 2007 |
| | TAPBP | Moderate | 1 | 0 | 55 | 2002 |
| | RGS9 | Moderate | 4 | 0 | 175 | 2004 |
| | HYAL1 | Moderate | 3 | 0 | 306 | 1999 |
| | EXOC2 | Limited | 1 | 100 | 12 | 2020 |
| | ZNF423 | Limited | 4 | 75 | 70 | 2012 |
| | RNASEH1 | Limited | 5 | 60 | 52 | 2019 |
| | CTF1 | Limited | 3 | 33 | 105 | 2000 |
| | BAZ2B | Limited | 9 | 33 | 22 | 2020 |
| | SARS1 | Limited | 3 | 33 | 36 | 2017 |
| | DNAJC7 | Limited | 10 | 30 | 24 | 2019 |
| | FBN2 | Limited | 72 | 28 | 303 | 1998 |
| | RHOH | Limited | 1 | 0 | 80 | 2012 |
| | IGKC | Limited | 3 | 0 | 112 | 1976 |
| | OTUD7A | Limited | 2 | 0 | 16 | 2020 |
| | ADCY1 | Limited | 1 | 0 | 89 | 2014 |
| | TOPBP1 | No Known | 2 | 50 | 250 | 2014 |
| | LRRC10 | No Known | 6 | 33 | 14 | 2015 |

| | | | | | | |
|---|---|---|---|---|---|---|
| | KMO | No Known | 1 | 0 | 160 | 2023 curation date |
| | CPT1B | No Known | 1 | 0 | 142 | 2021 curation date |
| **eval** | NLGN3 | Moderate | 54 | 35 | 2003 | 146 |
| | FOXE3 | Moderate | 24 | 29 | 2016 | 51 |
| | AHCY | Moderate | 18 | 28 | 2004 | 153 |
| | NDUFA2 | Moderate | 4 | 25 | 2008 | 36 |
| | TDO2 | Limited | 2 | 50 | 2017 | 173 |
| | PRPH | Limited | 10 | 20 | 2004 | 223 |
| | PTCD3 | Limited | 3 | 0 | 2019 | 10 |
| | KIF1B | No Known | 16 | 44 | 2001 | 188 |
| | PEX11G | No Known | 1 | 0 | 2020 | 7 |
| | MPST | No Known | 1 | 0 | 2023 curation date | 15 |

***Supplementary table 3: Curated dataset publication access and thresholding.*** *Across three evidence bases, we show the total and accessible papers considered for each gene after the GenCC ClinGen submission or curation date. Genes listed above NLGN3 were used in the dev set, while genes including and below NLGN3 were used in the eval set.*

| Evidence Base | Total Papers | Proportion Accessible (%) | # PubMed Papers |
|---|---|---|---|
| **Moderate (dev)** | 11.36 | 30 | 102.18 |
| **Limited (dev)** | 9.5 | 32.67 | 76.75 |
| **No Known (dev)** | 2.5 | 20.75 | 141.5 |
| **Moderate (eval)** | 25 | 29.25 | 96.5 |
| **Limited (eval)** | 5 | 23.33 | 135.33 |
| **No Known (eval)** | 6 | 14.67 | 70 |

***Supplementary table 4: Curated dataset average statistics per evidence base***. *Average number of PubMed papers, of which a proportion are accessible, and then filtered to those that are rare disease relevant (giving the total).*

Though less than 30% of publications would be accessible to EvAgg (open access papers with permissive licenses), the overall number of papers identified in the publication retrieval step was large enough that manually extracting content for all of them would have been intractable. Because of this, we randomly selected at most two papers for each gene from the subset of relevant papers in the PMC-OA dataset with licenses that permitted derivative use. Some of the 45 genes had only one or even zero relevant papers in this subset. See **Supplementary table 5** for a summary of the distributions of categorical values within the curated dataset.

| | dev M | dev L | dev N | eval M | eval L | eval N |
|---|---|---|---|---|---|---|
| paper build: hg19 | 11 | 15 | 1 | 36 | 1 | 1 |
| paper build: GRCh37 | 0 | 14 | 0 | 0 | 0 | 0 |
| paper build: GRCh38 | 2 | 0 | 0 | 0 | 0 | 0 |
| variant type: missense | 51 | 32 | 3 | 29 | 3 | 2 |
| variant type: splice region | 1 | 11 | 0 | 1 | 0 | 0 |
| variant type: stop gained | 17 | 9 | 0 | 8 | 0 | 0 |
| variant type: frameshift | 8 | 2 | 0 | 1 | 0 | 0 |
| variant type: frameshift insertion | 1 | 0 | 0 | 0 | 0 | 0 |
| variant type: inframe deletion | 0 | 1 | 0 | 0 | 0 | 0 |
| variant type: frameshift deletion | 2 | 1 | 0 | 0 | 0 | 0 |
| variant type: splice donor | 2 | 1 | 0 | 0 | 1 | 0 |
| variant type: synonymous | 2 | 1 | 0 | 0 | 0 | 0 |
| variant type: indel | 1 | 0 | 0 | 0 | 0 | 0 |
| variant type: stop lost | 0 | 0 | 0 | 9 | 0 | 0 |
| zygosity: het. | 14 | 19 | 3 | 9 | 3 | 2 |
| zygosity: homo. | 4 | 6 | 0 | 27 | 1 | 0 |
| zygosity: compound het. | 9 | 4 | 0 | 8 | 0 | 0 |
| var. inherit.: maternal | 3 | 10 | 0 | 0 | 0 | 0 |
| var. inherit.: paternal | 3 | 4 | 0 | 0 | 1 | 0 |
| var. inherit.: mat. & pat. homo. | 2 | 2 | 0 | 0 | 0 | 0 |
| var. inherit.: de novo | 4 | 2 | 0 | 0 | 0 | 0 |
| license: CC BY | 79 | 39 | 3 | 9 | 4 | 2 |
| license: none | 0 | 12 | 0 | 0 | 0 | 0 |
| license: CC0 | 4 | 0 | 0 | 8 | 0 | 0 |
| license: CC BY-NC-SA | 0 | 6 | 0 | 0 | 0 | 0 |
| license: CC BY-NC | 2 | 1 | 0 | 31 | 0 | 0 |
| individual observed cases per paper: mean | 0.93 | 3.15 | 0.5 | 7.33 | 1 | 1 |
| individual observed cases per paper: median | 1 | 2 | 0.5 | 2.5 | 1 | 1 |
| individual observed cases per paper: mode | 0 | 2 | 0 | 0 | 0 | 1 |
| # unique papers | 15 | 13 | 2 | 6 | 3 | 2 |

***Supplementary table 5: Curated dataset statistics**. M, moderate evidence type; L, limited evidence type; N, no known gene-disease relationship type.*

##### *Curated dataset generation rubric*

Please note that this rubric was generated by and for our internal curation team, and it was accompanied by additional framing, requirements, clarifying comments and notes which are not shared here. The following text was not modified for this publication. As such, some terminology differs from the main text, including individual observations (referred to as “patients” here), error analysis (referred to as “discrepancy resolution” here), paper finding (referred to as “gathering the right papers” and “picking papers” here), and content extraction (referred to as “gathering the right content” here).

**RUBRIC** (red - added in response to discrepancy resolution task)

- Gathering the right papers, towards (1):
  - Include
    - Variant (SNV/ InDels) reported in a human with a *reasonably* monogenic condition
    - Functional papers that model patient variants
    - Review papers that only report previously reported variants
    - Papers that refer to a previously reported variant in the same family, but with extra data.
    - Papers that are behind paywalls but we can clearly tell they fit the criteria from the abstract
    - Syndromic, inherited cancer
    - Include benign/VUS reports
  - Exclude for v1 (highlight in yellow)
    - Digenic, familial
    - Only report of a variant in the paper is an SV
    - Animal model about a gene but no specific human equivalent mutation
    - Not in English
    - the paper's **abstract** not accessible (e.g. PMC OA with non-ND license)
    - Papers where we can't check if it is a right paper or not (pay wall).
    - Not somatic cancer
  - Filters in PubMed
    - Start with ClinGen's submission to GenCC's start date

- Gathering the right content in those papers, towards (2):
  - When picking papers:
    - Do not include content from the supplement
    - the paper must be accessible (PMC OA with non-ND license only)
      - Can we extract enough information from abstracts? This can be discussed during development
    - The papers must be scannable (e.g. not too old)
    - the variants in the paper should be accessible in the main paper
      - Or mention that is it first mentioned in supplement
    - the papers should be case report, case series or cohort analysis
      - we may not have variants reported in cohort analysis
      - *Case report*: one patient report
      - *Case series*: multiple patients with variants in a single gene
      - *Cohort analysis*: site-wide assessment of rare disease patients
    - there should be variants listed in the paper

#### Pipeline evaluation

Before finalizing the evaluation set, we took the opportunity to correct obvious errors (e.g., typos) in the evaluation curated dataset that were uncovered during pipeline evaluation execution.

For paper selection we calculated recall as the proportion of manually curated papers that EvAgg retrieved, and precision as the proportion of papers returned by EvAgg that were included in the curated dataset (see equations below). Note that these comparisons were made after excluding all papers from the curated dataset that were inaccessible to EvAgg. Inaccessible papers were those that were not included in PMC-OA, those that had licenses that did not permit derivative use, or those that were not returned based on a date-restricted search for the gene on the PubMed website.

$$precision = \frac{correct\ \#\ retrieved\ [task\ variable]}{total\ \#\ retrieved\ [task\ variable]}$$

$$recall = \frac{correct\ \#\ retrieved\ [task\ variable]}{total\ \#\ correct\ [task\ variable]\ in\ truth\ set}$$

For observation finding, recall was calculated as the proportion of curated dataset observations retrieved and precision was the proportion of observations returned by EvAgg that were included in the curated dataset. Each unique observation was identified by a variant-individual pair. Note that to account for slight variations in LLM outputs and manual curation of observations, the following individual identifiers were treated as equivalent during pipeline evaluation: "the proband", "proband", "the patient", "patient", "one proband", and “unknown”. To better understand whether missing or extraneous observations could be attributed to entity recognition (i.e., finding the correct variants within a paper) or entity linking (correctly associating those variants with individuals), we separately assessed precision and recall of variant finding using the same approach while treating all individual identifiers as equivalent.

It is important to note that because the pipeline involves successive stages of processing (paper selection -> observation finding -> content extraction), in specific runs it is possible that different sets of papers or different observations found in those papers will impact the inputs to and corresponding outputs of subsequent stages. Put another way, because of inherent variability in LLM responses, the exact same papers are not being processed across each of these five runs. To better understand the impact of LLM variability, the results section includes an assessment of the between-run consistency for paper finding.

Finally, for all content categories except phenotype, we assessed accuracy by comparing the proportion of manually curated content that exactly matched the values produced by EvAgg.

$$accuracy = \frac{\#\ correct\ content\ values}{total\ \#\ content\ values\ in\ truth\ set}$$

##### *Phenotype extraction pipeline evaluation*

When performing pipeline evaluation on phenotype extraction, we attempted to assess whether the pipeline was capable of identifying all relevant phenotypes possessed by a patient, without overly penalizing for subtle errors. To accomplish this, we imposed hierarchical reasoning to generalize specific phenotype terms from publication text to organ/system description phenotype terms. Organ/system description HPO terms were defined as all terms where the shortest path from that term to the HPO root entity (HP:0000001 | All) is exactly 2 steps (see **Supplementary table 1** for a list of all organ/system level terms considered for pipeline evaluation of phenotype extraction performance). For each specific HPO term we identified the organ/system description level term closest (via path length) using the “path_to_other” function in the pyhpo Python library. Using these generalized representations of an observation’s corresponding phenotype, we could then assess whether all generalized phenotypes present in the curated dataset were also present in the pipeline's output and whether the pipeline extracted any "extra" generalized phenotypes.

Although EvAgg does not constrain phenotype extraction outputs beyond ensuring HPO terms are valid, every phenotypic term in the curated dataset generalizes to organ/system description child terms of parent term "HP:0000118 | Phenotypic abnormality" (see **Supplementary table 1** for full list) as do the vast majority of pipeline outputs (~98% across multiple pipeline evaluation runs). Around two percent of pipeline outputs generalize to a different child of "HP:0000001 | All" (i.e. children of "HP:0000005 | Mode of inheritance", "HP:0012823 | Clinical modifier", "HP:0020228 | Biospecimen phenotypic feature", 'HP:0032223 | Blood group", "HP:0032443 | Past medical history", or "HP:0040279 | Frequency").

To evaluate phenotype extraction performance of the EvAgg pipeline, we asked whether each of these generalized terms that were present in the curated dataset were also present in pipeline output and vice versa, calculating accuracy, recall, and precision of phenotypic descriptions. Accuracy was defined as the percentage of observations for which the exact set of generalized organ/system description phenotypic terms from the curated dataset was present in the pipeline output, with zero extraneous terms. Recall was the percentage of observations for which all generalized phenotypic terms from the curated dataset were present in pipeline outputs, regardless of extraneous terms. Precision was the percentage of observations for which there were zero extraneous terms, regardless of the proportion of correct terms found. By computing this set of metrics, we could develop a complete understanding of the pipeline's ability to extract observed phenotypes and the typical failure modes leading to reduced accuracy.

Because the HPO is a directed-acyclic graph and not a hierarchy, it is possible that a given HPO term extracted from a paper's text would have multiple ancestors at the organ/system description level. Indeed this was the case for approximately 25% of the unique terms extracted during pipeline evaluation analyses. However, for simplicity, the only organ/system description level ancestor terms considered during our primary pipeline evaluation analyses were those with the shortest path within the HPO to the extracted term.

We evaluated an alternative phenotype approach to account for the possibility of multiple organ/system description generalization terms, however this had no substantial impact on our findings. For this "multiple organ/system description generalizations" phenotype pipeline evaluation, we considered a phenotype term a match if any of the organ/system description terms corresponding to a specific EvAgg pipeline output term matched with any of the organ/system description terms corresponding to a curated dataset term (as opposed to the consideration of a single organ/system description parent term when multiple were possible). This didn't have a substantial impact on phenotype evaluation performance: eval set accuracy went from 0.862 with single shortest path organ/system description term to 0.878 when considering a match with any parent term when multiple generalizations were possible, recall was the same (0.961), and precision increased from 0.892 to 0.908.

##### *Error analysis*

After executing the five replicates of pipeline runs on the development and evaluation gene sets, we compiled consistent disagreements between pipeline outputs and the curated dataset. Errors were defined as publications in the curated dataset not retrieved by EvAgg or publications retrieved by EvAgg which were not in the curated dataset in 3 or more runs, observations in the curated dataset not found by EvAgg or observations found by EvAgg not in the curated dataset in 3 or more runs, or extracted content where pipeline outputs were discordant from the curated dataset half of the time or more. For phenotypes, an entire observation was categorized as disagreeing with the curated dataset if any of the generalized phenotypes above were consistently discordant. We only considered content discrepancies where at least 2 of the 5 execution replicates processed that paper/observation during content extraction.

The output of this review process was analyzed separately to assess consistent, true error modes in pipeline output and the extent to which the EvAgg pipeline was able to identify additional relevant information in the literature that was not included during initial generation of the curated dataset. Furthermore, after error analysis completion, warranted updates were made to a revised version of the curated dataset. We present the error analysis review in the main text as well as EvAgg pipeline performance on both the original and revised curated datasets below (**Supplementary table 6**).

#### User study design and analysis

User study design underwent subject matter expert review and IRB review (OHRP-IRB00009672) at Microsoft.

To minimize the impact of differences between study participants and the difficulty of individual cases, we created fixed-order, approximately difficulty-matched sets of four unsolved cases each selected from the Rare Genomes Project (raregenomes.org). Difficulty was determined by quantity of variants returned in fixed variant "standard" searches within *seqr* (de novo/dominant, recessive, restrictive and permissive), ClinGen/GenCC annotations of genes returned in search, and affected status of individuals within family pedigree. Evidence Aggregator results were precomputed and made available for genes returned in variant search results that also had a ClinGen annotation value of moderate, limited, or no-known gene-disease relationship (per pipeline evaluation criteria). User study participants were randomized to case analysis with either case set A or B on the first day, analyzing the reciprocal set on the second day.

We collected user experience data from Likert-scale surveys and semi-structured interviews. In surveys following use of EvAgg, we solicited user experience information to compare user experience during case analysis with/without the EvAgg outputs, as well as user sentiment and aspects of trust towards AI-derived evidence. Discussions solicited additional feedback about specific features contributing to sentiment and trust, importance of accuracy, characterization of how EvAgg may impact their workflow, and suggested changes which would improve their user experience.

Quantitative data were collected via time-stamped user action recording during case review, as well as timestamped artifacts generated by interactions with *seqr* (custom variant searches, tagging variants, and taking notes). Timestamped user actions were collected in real time during study sessions and were verified via video recording review and comparison to *seqr* timestamps to ensure accuracy ±2 seconds. The user actions for which timestamps were recorded included: starting to review a case, starting to review a new variant, starting to re-review a saved variant, completing review of a variant (new or saved), performing a search in *seqr*, searching for publications using external tools, and publication reading. Analysts were asked to work at their normal pace, and cases were considered "complete" when they stated they were "done" analyzing a case and would normally move on (study administrator would verify with participants that they considered their case analysis complete prior to moving on to the next case). The number of variants reviewed was defined by "opening up" a different resource to investigate variant further (i.e. opening the EvAgg table, clicking the gene name to open a pop-up in *seqr* with additional gene information, navigating to OMIM/Decipher, etc.). Publication reading was defined as opening up a publication's full text (usually via PubMed Central or individual publisher websites) and spending time reviewing the text (i.e. reading the abstract, skimming through text, thoroughly reading sections with individual case information, searching text for keywords to find information of interest and reading highlighted areas, etc.).

We used a linear mixed-effects model to analyze the impact of EvAgg on individual behavioral metrics, accounting for both fixed effects (study session and case set) and random effects (individual participants) under the assumption that there could be large inter-individual differences in case review approaches. All statistical modeling was carried out using the `mixedlm` function from the `statsmodels` python package.

### Supplementary results

#### Error analysis results

**Supplementary table 6** details results from manual assessment of putative EvAgg errors as compared to the curated dataset. The curated dataset was updated when putative EvAgg errors were ultimately deemed to be correct by the curation team.

| Task | EvAgg and curated dataset agreement | Total disagreements (putative errors) | Curated dataset updated (%) | EvAgg errors (%) | EvAgg missed – False negative (%) | EvAgg extras – False positive (%) |
|---|---|---|---|---|---|---|
| All | 1423 | 470 | 235 (50%) | 235 (50%) | - | - |
| Paper selection | 104 | 74 | 52 (70%) | 22 (30%) | 3 (5%) | 19 (26%) |
| Observation finding | 241 | 81 | 49 (60%) | 32 (40%) | 8 (10%) | 24 (30%) |
| Phenotype | 702 | 226 | 96 (42%) | 130 (58%) | - | - |
| V. inheritance | 122 | 45 | 17 (38%) | 28 (62%) | - | - |
| V. type | 118 | 13 | 2 (15%) | 11 (85%) | - | - |
| V. zygosity | 136 | 31 | 19 (61%) | 12 (39%) | - | - |

***Supplementary table 6: Error analysis results.*** *After an initial pipeline evaluation run, putative errors were manually assessed to determine EvAgg error modes and refine curated dataset for pipeline evaluation and curated dataset release (see Data and Code Availability). Percent is calculated as percent of total disagreements per task. Performance for EvAgg was optimized to favor recall over precision, i.e. optimizing for low false negative rate rather than low false positive rate. EvAgg, Evidence Aggregator; V., variant.*

#### Full pre- and post-error analysis pipeline evaluation performance

**Supplementary table 7** provides complete detail as to pipeline evaluation performance on both the dev and eval gene sets before and after error analysis.

| Task | Set | Pre-error-analysis performance | | | Post-error-analysis performance | | |
|---|---|---|---|---|---|---|---|
| | | Precision (std) | Recall (std) | Accuracy (std) | Precision (std) | Recall (std) | Accuracy (std) |
| **Paper selection** | dev | 0.658 (0.017) | 0.797 (0.012) | - | 0.882 (0.016) | 0.971 (0.006) | - |
| | eval | 0.563 (0.019) | 0.916 (0.012) | - | 0.832 (0.028) | 0.970 (0.017) | - |
| **Observation finding** | dev | 0.716 (0.029) | 0.888 (0.026) | - | 0.829 (0.028) | 0.920 (0.02) | - |
| | eval | 0.624 (0.037) | 0.746 (0.058) | - | 0.769 (0.042) | 0.784 (0.055) | - |
| **Variant Finding** | dev | 0.805 (0.022) | 0.934 (0.008) | - | 0.883 (0.021) | 0.950 (0.009) | - |
| | eval | 0.666 (0.026) | 0.958 (0.000) | - | 0.821 (0.018) | 0.979 (0.019) | - |
| **Content extraction: Phenotype** | dev | 0.691 (0.045) | 0.874 (0.034) | 0.628 (0.033) | 0.745 (0.042) | 0.0886 (0.015) | 0.659 (0.025) |
| | eval | 0.810 (0.036) | 0.932 (0.029) | 0.798 (0.050) | 0.891 (0.033) | 0.961 (0.025) | 0.862 (0.047) |
| **Content extraction: Zygosity** | dev | - | - | 0.785 (0.021) | - | - | 0.800 (0.022) |
| | eval | - | - | 0.934 (0.036) | - | - | 0.922 (0.027) |
| **Content extraction: Variant type** | dev | - | - | 0.837 (0.102) | - | - | 0.841 (0.092) |
| | eval | - | - | 0.844 (0.014) | - | - | 0.888 (0.012) |
| **Content extraction: Inheritance** | dev | - | - | 0.808 (0.013) | - | - | 0.757 (0.013) |
| | eval | - | - | 0.520 (0.074) | - | - | 0.768 (0.076) |

***Supplementary table 7: EvAgg pipeline evaluation performance pre- and post-error analysis.*** *All values reported are the average (mean) across the 5 repeated tool executions for the dev and eval gene sets. STD, standard deviation.*

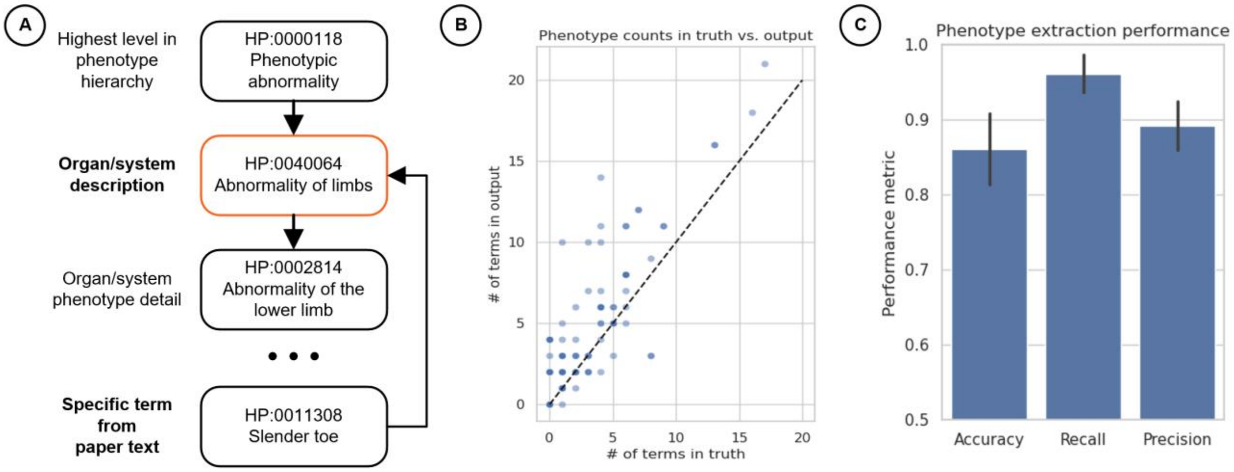


***Supplementary figure 2: Pipeline evaluation of phenotype methodology and results.*** *(A) Depiction of the phenotype generalization approach. Specific HPO terms derived from paper text were generalized to corresponding organ/system description terms in the HPO (generally a child of parent term "HP:0000118 – Phenotypic abnormality") before making comparisons. (B) This scatter plot shows that EvAgg consistently identifies more phenotypic terms per observation than are found in the curated dataset. (C) Bar plot separately showing the accuracy, recall, and precision of phenotype extraction. Accuracy is defined as the proportion of observations with a perfect match of generalized organ/system description terms, recall defined as the proportion of observations where all curated dataset organ/system description terms were included in EvAgg's output, and precision defined as the proportion of observations where there were no extraneous terms included in EvAgg's output. Relatively high recall as compared to precision is concordant with the consistent overidentification of HPO terms shown in subplot B.*

#### Pipeline performance LLM comparison

While the landscape of publicly accessible LLMs, their available capabilities, and associated costs is rapidly changing, it is still helpful to observe a comparison snapshot of the current set of models available on AOAI that met pipeline requirements at the time of analysis. Models considered in this analysis needed to support at least 128,000 input tokens and provide responses in validated JSON format. At the time of authoring this manuscript the models available that met these requirements were GPT-4 Turbo (1106-Preview), GPT-4o (2024-08-6), and GPT-4o-mini (2024-07-18). While GPT-4-Turbo and GPT-4o are both estimated to have 1.76T parameters, GPT-4o-mini is more than two orders of magnitude smaller with 8B parameters. The latter model is correspondingly much cheaper to run, with retail prices on the Azure OpenAI Service at least an order of magnitude lower than its larger counterparts.

Cost may be a factor when considering use of EvAgg for genes with a definitive or strong GDR, as these genes would be expected to have significantly more publications to process than genes with moderate/limited/no-known gene disease relationships. The inclusion of a new module or LLM prompt to modify functionality (e.g. variant-centric search for strong/definitive GDR genes or to leverage user credentials to obtain access to closed-access journals they have appropriate access to) would also increase cost. Future work would be required to explore such modified applications.

The EvAgg pipeline was configured to run using each of these models and both the development and evaluation gene sets. Each of these configurations was executed 5 times to provide more accurate estimates of performance. The same pipeline evaluation methodology as described elsewhere was applied to the outputs of these runs, comparing them to the updated curated dataset. One run for GPT-4o on the test set was discarded from downstream analysis due to a technical issue (output token generation overrun). Performance for this run was imputed as the mean of the other 4 runs for this condition for statistical analyses.

While the absolute differences between model performance on the paper selection task was not large, GPT-4-Turbo significantly outperformed the other two models in terms of recall on the paper selection task ($p < 0.05$ via 2-way repeated measures ANOVA with Tukey's HSD for within factor comparisons throughout). The performance difference between the other two models on this task was not significant. However, for the observation finding task recall was substantially lower using GPT-4o-mini, particularly on the test set where average recall dropped to 0.173. Both GPT-4-Turbo and GPT-4o significantly outperformed GPT-4o-mini on this task ($p < 0.05$) but were not significantly different themselves. Taken together these results suggest that for certain applications, smaller models may perform sufficiently well, but as task complexity and relevant context increase, larger models currently outperform smaller ones.

Because pipeline evaluation for downstream content extraction tasks (e.g., determining the phenotype of an affected individual) depend heavily on the specific observations found by the pipeline and the performance on observation finding was so disparate, a direct comparison of LLM model performance on content fields is not feasible.

See **Supplementary table 8** and **Supplementary figure 3** for a detailed comparison of EvAgg pipeline evaluation based on different underlying LLMs.

| **Model** | **Set** | **Paper selection** | | | **Observation finding** | | **Variant finding** | |
|---|---|---|---|---|---|---|---|---|
| | | Runtime (std) | Precision (std) | Recall (std) | Precision (std) | Recall (std) | Precision (std) | Recall (std) |
| **GPT-4-Turbo** | dev | 13900 (363) | 0.882 (0.016) | 0.971 (0.006) | 0.829 (0.028) | 0.920 (0.020) | 0.883 (0.021) | 0.950 (0.009) |
| | eval | 9160 (380) | 0.832 (0.028) | 0.970 (0.017) | 0.769 (0.042) | 0.784 (0.055) | 0.821 (0.018) | 0.979 (0.019) |
| **GPT-4o** | dev | 2660 (120) | 0.884 (0.022) | 0.948 (0.013) | 0.736 (0.032) | 0.910 (0.038) | 0.826 (0.026) | 0.944 (0.023) |
| | eval | 1550 (45.8) | 0.832 (0.017) | 0.932 (0.022) | 0.672 (0.128) | 0.594 (0.195) | 0.845 (0.031) | 0.931 (0.040) |
| **GPT-4o-mini** | dev | 2470 (200) | 0.776 (0.036) | 0.948 (0.009) | 0.673 (0.026) | 0.786 (0.044) | 0.779 (0.023) | 0.902 (0.023) |
| | eval | 1500 (135) | 0.784 (0.030) | 0.898 (0.029) | 0.309 (0.096) | 0.173 (0.036) | 0.707 (0.041) | 0.833 (0.088) |

***Supplementary table 8: Paper selection and observation finding pipeline performance model comparison.*** *Precision and recall are as defined above. Paper selection runtime is the average (mean) duration of pipeline execution for the relevant gene set (dev or eval), provided in seconds. STD, standard deviation. Note the catastrophic reduction in pipeline performance on observation finding for GPT-4o-mini relative to the other models. STD, standard deviation.*

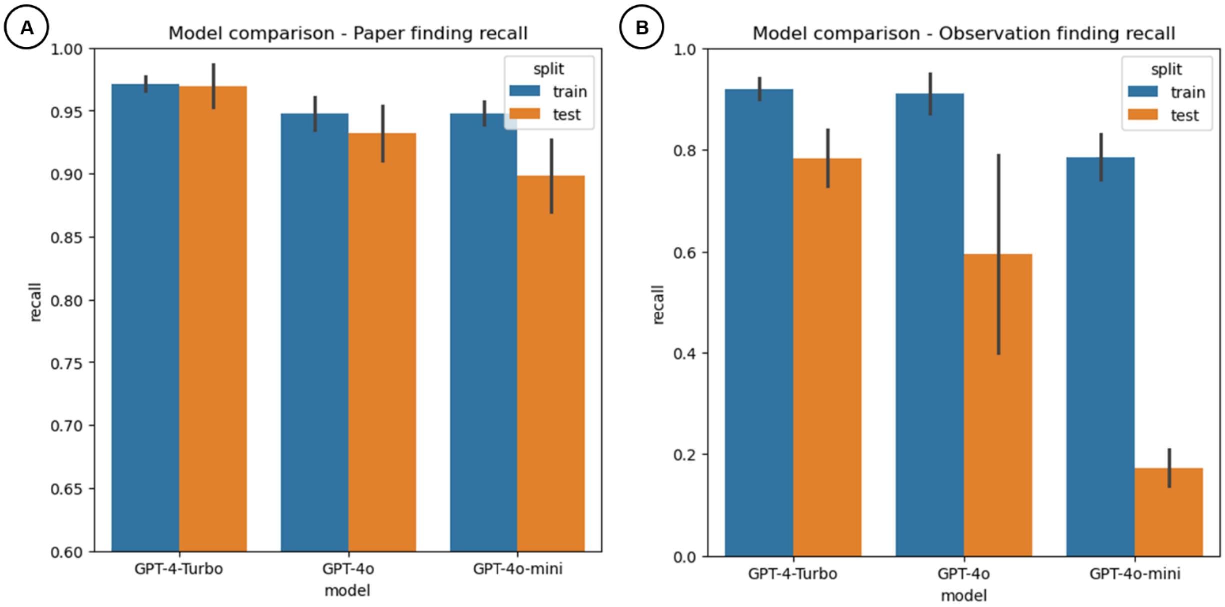


***Supplementary figure 3: Pipeline performance model comparison.*** *Comparison of the three models with sufficient input context length to support EvAgg (128k tokens). Note fairly consistent performance across the three models for the task of paper finding (A), and substantially poorer performance on observation finding (B) for GPT-4o-mini.*

#### User study analysis results

See **Supplementary table 9** below for demographic information on user study participants. See **Supplementary Figure 4** for additional user experience impressions.

| | N | % |
|---|---|---|
| **All participants** | 8 | |
| **Gender** | | |
| Woman | 6 | 75.0 |
| Man | 2 | 25.0 |
| **Ethnicity/Descent** | | |
| Caucasian/European | 4 | 50.0 |
| African | 2 | 25.0 |
| Latin/Hispanic/Spanish | 2 | 25.0 |
| South Asian | 1 | 12.5 |
| **Education/certifications** | | |
| MD | 4 | 50.0 |
| PhD | 7 | 87.5 |
| CGC | 1 | 12.5 |
| LCGC | 1 | 12.5 |
| MS | 1 | 12.5 |
| **Rare disease case analysis experience** | | |
| 1-5 years | 3 | 37.5 |
| 5-10 years | 5 | 62.5 |
| **Experience using *seqr*** | | |
| Less than 6 months | 1 | 12.5 |
| 1-2 years | 3 | 37.5 |
| 3-5 years | 1 | 12.5 |
| More than 5 years | 3 | 37.5 |
| ***seqr* access method** | | |
| Organization *seqr* instance | 7 | 87.5 |
| Terra | 1 | 12.5 |
| **Work organization type** | | |
| Academic research institute | 7 | 87.5 |
| Hospital/medical centers | 1 | 12.5 |

***Supplementary table 9*: *User study participant demographics.*** *MD, Doctor of Medicine; PhD, Doctor of Philosophy; CGC, Certified Genetic Counselor; LCGC, Licensed Certified Genetic Counselor; MS, Master of Science*

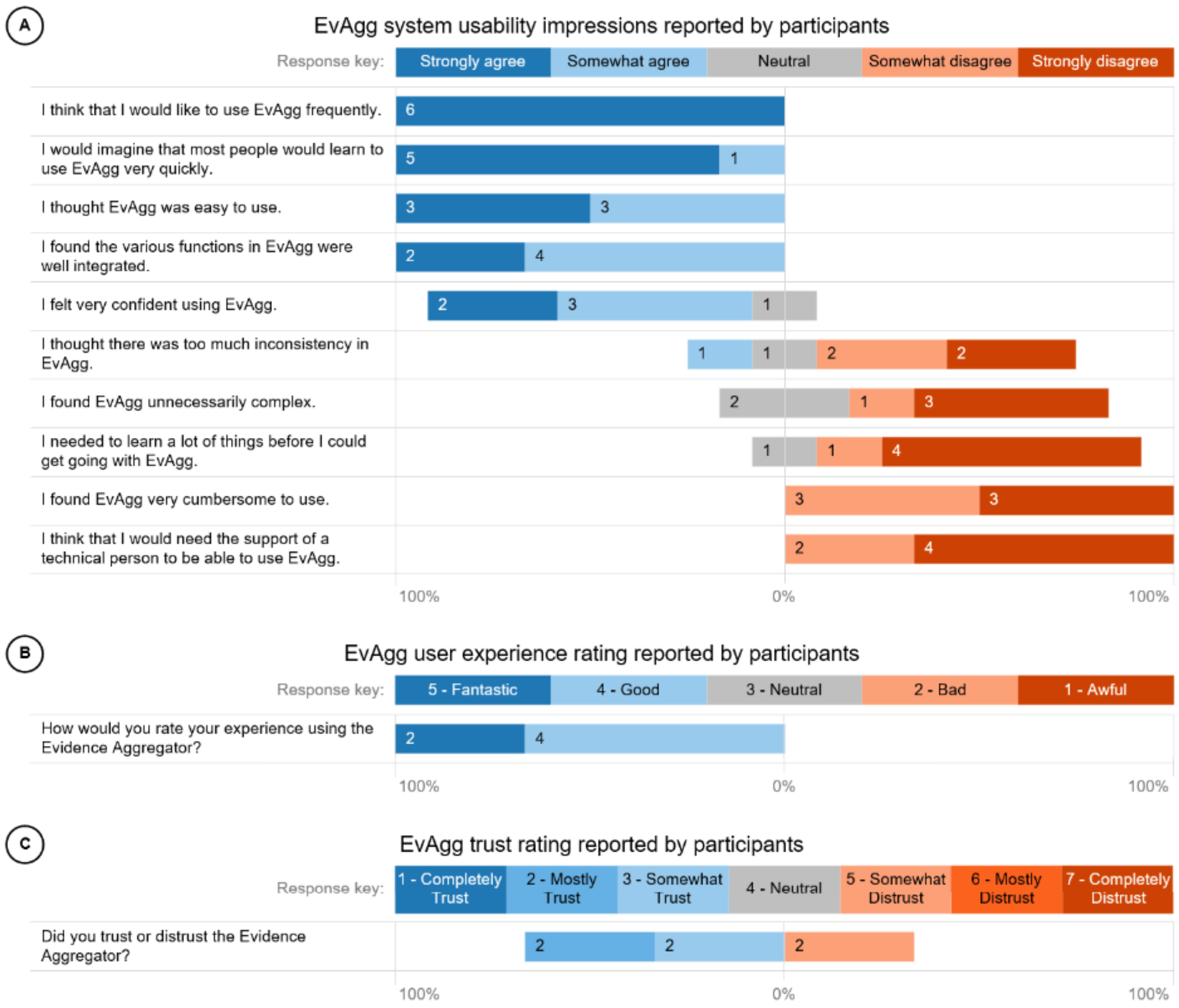


***Supplementary figure 4:*** *Additional user experience impressions as reported by participants (A) System usability impressions on a 5-point Likert scale. Agreement with statements ranged from "Strongly agree" to "Strongly disagree", from system usability score (SUS) rating system. (B) User sentiment from survey data reported on a 5-point Likert scale: scores ranged from 1 (strongly dislike: "Awful") to 5 (strongly like: "Fantastic"). (C) User trust from survey data reported on a 7-point Likert scale: scores ranged from 1 ("completely trust") to 7 ("completely distrust"). Response count is reported within each stacked bar segment.*

##### *Subsequent Rare Genomes Project variant review*

A subset of variants tagged for review were further evaluated by Rare Genomes Project expert variant analysts at the Broad Institute of MIT and Harvard: variants tagged for review by at least 3 analysts (from either study session) and variants only tagged for review by multiple analysts when EvAgg was available for the variant (which would mean 0/4 analysts tagged the variant for review during the first baseline session, and at least 2/4 analysts tagged the variant for review when they had the use of EvAgg). While no diagnoses have resulted from the user study, two variants were submitted to Matchmaker Exchange: one variant tagged for review by 7/8 study participants and one variant which was only tagged for review when EvAgg was available.

##### *Case analysis behavior*

Full user study survey and timestamped workflow data are available in the GitHub repository (see **Data and Code Availability**). See **Supplementary Figure 5** for user study measures per study session stratified by case set and participant.

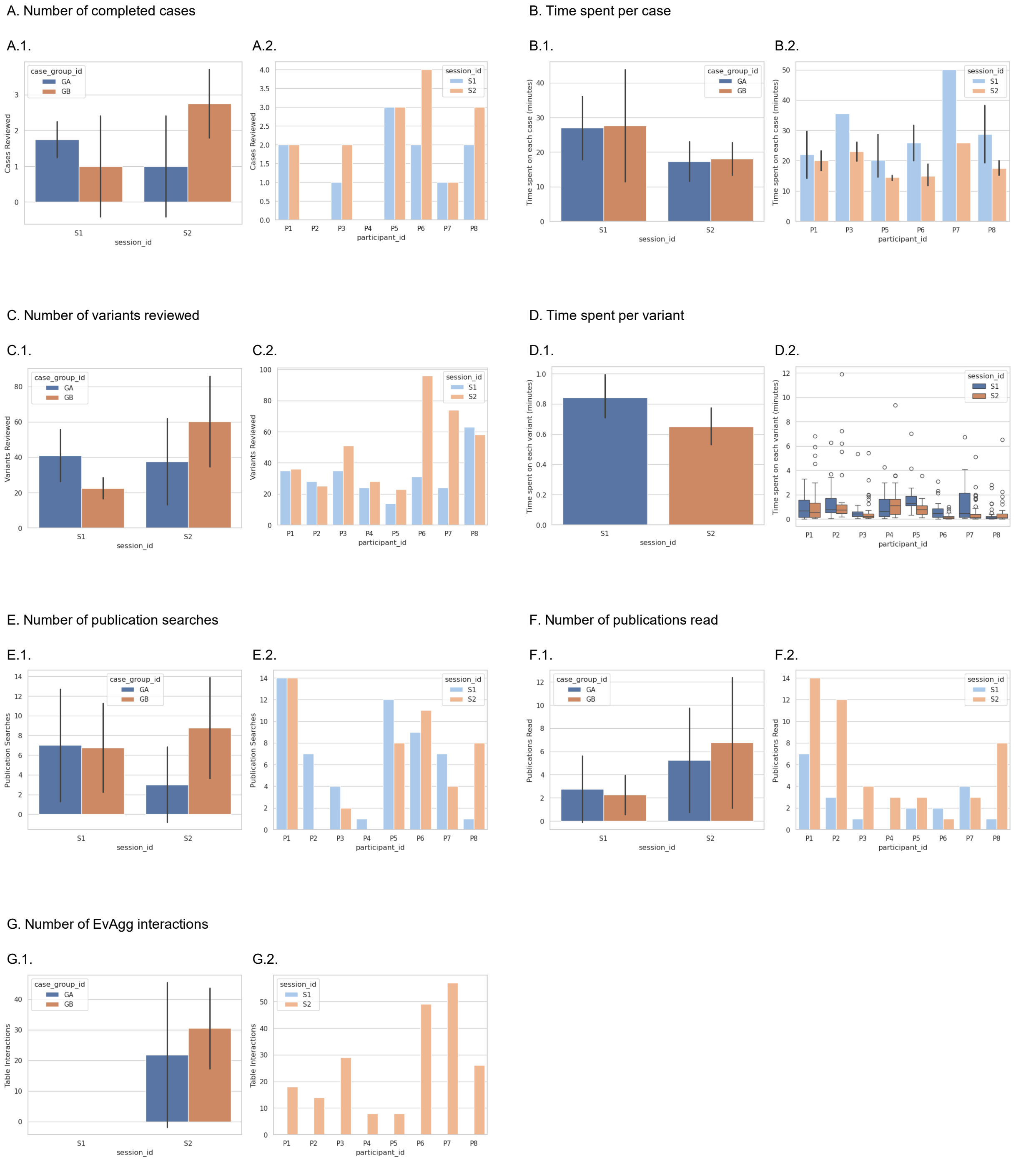


***Supplementary figure 5: User study measures per study session stratified by case set and participant.*** *The user study consisted of two 1-hour sessions of case analysis by expert analysts, with session 1 serving as a baseline and session 2 with use of EvAgg. Reported measures: (A) number of completed cases, (B) time spent per case (completed cases only), (C) number of variants reviewed, (D) time spent per variant, (E) number of publication searches, (F) number of publications read, (G) number of EvAgg interactions. For each measure: the bar plot on the left (1) shows the average value and standard deviation per*

*session (session_id; S1= session 1, S2= session 2) subdivided by case set (case_group_id; GA in blue= case set A, GB in orange = case set B); the bar plot on the right (2) shows measure values for each participant subdivided by study session, time spent per case (B.2.) displays average and standard deviation, time spent per variant (D.2.) is displayed as a box-and-whisker plot to better visualize the distribution of time spent on each variant as well as highlight outliers. This figure is intended to inform the reader of observed workflow measure differences between case sets and among participants.*

Two participants did not complete a full case analysis during their 1 hour of case analysis, meaning those study sessions are not included in the analysis of time spent per case. Cases that were incomplete at the end of the 1 hour of case analysis were also not included in the analysis of time spent per case. As EvAgg was unavailable during the first study session, there are no results reported for number of EvAgg table interactions for that session. Notably, while the number of variants reviewed increased between the two sessions (31.8 ± 14.4 vs 48.9 ± 26.1), this change was not significant.
